# High-throughput Approaches to Uncover Synergistic Drug Combinations in Leukemia

**DOI:** 10.1101/2022.11.29.518409

**Authors:** Emma J. Chory, Meng Wang, Michele Ceribelli, Aleksandra M Michalowska, Stefan Golas, Erin Beck, Carleen Klumpp-Thomas, Lu Chen, Crystal McKnight, Zina Itkin, Sanjay Divakaran, James Bradner, Javed Khan, Berkley E. Gryder, Craig J. Thomas, Benjamin Z. Stanton

## Abstract

We report a comprehensive drug synergy study in acute myeloid leukemia (AML). In this work, we investigate 11 cell lines spanning both MLL-rearranged and non-rearranged subtypes. The work comprises a resource for the community, with many synergistic drug combinations that could not have been predicted *a priori*, and open source code for automation and analyses. We base our definitions of drug synergy on the Chou-Talalay method, which is useful for visualizations of synergy experiments in isobolograms, and median-effects plots, among other representations. Our key findings include drug synergies affecting the chromatin state, specifically in the context of regulation of the modification state of histone H3 lysine-27. We report open source high throughput methodology such that multidimensional drug screening can be accomplished with equipment that is accessible to most laboratories. This study will enable preclinical investigation of new drug combinations in a lethal blood cancer, with data analysis and automation workflows freely available to the community.

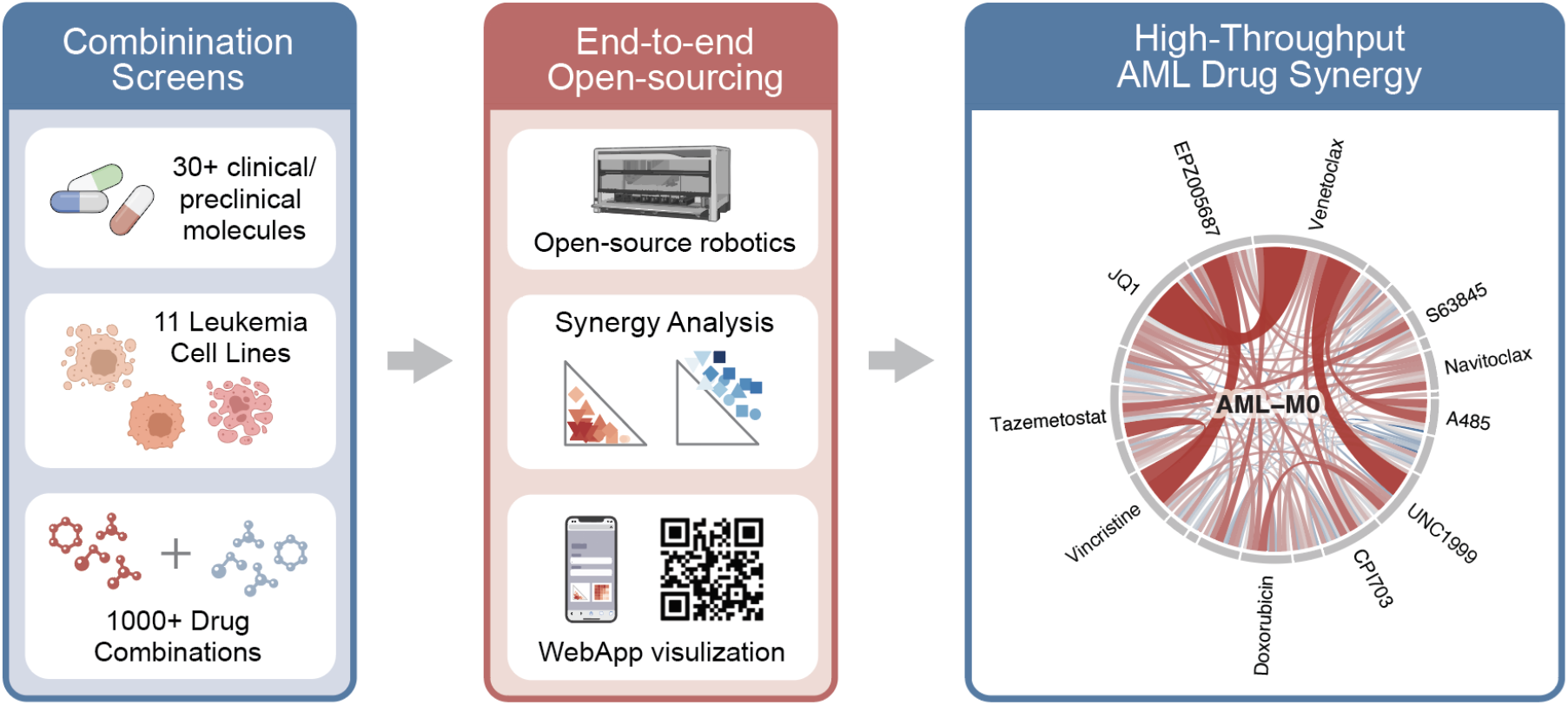

## INTRODUCTION

The local chromatin environment is essential for generating the context in which genes are repressed or activated. However, genetic insertions and deletions, as well as changes to the epigenome can generate new context-specific patterns of transcription ^1–3^. Thus, if the local context for gene expression can be epigenetically modified, this presents a paradigm where the functional targeting of gene expression with epigenetic probes, inhibitors, and drugs can also be modified. In our studies, we sought to modify the functional relevance of gene-drug interactions through the systematic introduction of secondary agents which we hypothesized would potentiate or antagonize the effects of chromatin-targeting molecules.

In this work, we investigate the functional interactivity of molecules targeting the following areas: (1) histone acetylation, (2) histone methylation, (3) chromatin reading, (4) cytoskeletal, (5) DNA replication, and (6) apoptosis, and selected our candidates for combination studies from (1) clinical, (2) pre-clinical and (3) basic science focused probe compounds. The strategy for combining two drugs for enhanced potency ^4^ has been broadly successful and benefited from recent genomic and machine learning-guided approaches to identify and predict drug interaction networks ^5,6^. Thus, we sought to uncover functional drug interactions analogous to genetic mutations that can enhance or suppress phenotypes similar to second-site suppressor (or activator) mutations from genetic screens, in a high-throughput manner in line with these efforts. Our approach for uncovering phenotypic drug interactions in leukemia allowed for the assessment of hundreds of pairwise comparisons rapidly, with many candidates that already have proceeded through clinical testing. Our approach for miniaturized combination phenotypic screening for enhancers or suppressors of leukemia viability in high-throughput is highly modular and allowed us to compare drug interactivity and reproducibility across each compound class in our system. To enhance the accessibility of this screening platform, we implemented an automated Chou-Talalay Synergy R-based analysis method ^7,8^, and additionally developed an open-source automation method using PyHamilton for liquid-handling robots ^9^. Collectively, this work greatly increases the ease-of-adoptability for scientists and clinicians seeking to identify novel combinations of clinically promising drug synergy, potentiation, or to identify drug antagonism in cases where *a priori* contraindication has not been thoroughly examined.

The work presented herein includes the rational design of high-throughput drug combinations for rapid testing, a new open-source analysis pipeline for evaluation of drug synergy and antagonism, and basic computational resources for establishing drug synergy experiments without necessitating a fully-automated operation automation screening facility, rather a single liquid handling robot. We emphasize compound efficacy in both single-agent and combination contexts, as part of our study design. Ultimately our experimental readout was functional, with cell viability measurements after 2-day single agent or combination drug incubations. As we discuss, this experimental strategy was especially useful when evaluating potentiation of epigenetic drugs altering histone H3 lys-27 methylation in AML, as EZH2 single-agent inhibition generally had minimal effects on cell viability within a two-day period. Our study highlights the utility of epigenetic drug combinations using the context of AML, especially where effects on chromatin structure that may be non-toxic to tumors as single agent effects can be strongly potentiated with secondary agents.

## MATERIALS AND METHODS

### Combination Synergy Screening with the Chou-Talalay Method

The concept of additivity in drug interactions is key to defining molecular synergy ^10^. We can define additive interactions in the context of the *fraction unaffected* (*f*_*u*_). If a combination of drugs, A and B, is additive, the *fraction unaffected* by treatment of the combination (*f*_*u*_)_A+B_ will be equal to the product of (*f*_*u*_)_A_ x (*f*_*u*_)_B_ over the same incubation time period ^11^. Thus, a synergistic drug combination has the property:

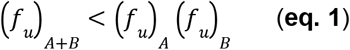

In our studies, the fraction unaffected equates to the fraction of viable fraction of leukemia cells after 2-day drug treatments, which can be quantified rapidly in high-throughput with viability assays.

Likewise, antagonism can be expressed such that the (*f*_*u*_)_A+B_ value is greater than the fraction unaffected for either drug as a single agent. The ratio of the *fraction affected* (*f*_*a*_) with the *fraction unaffected* (*f*_*u*_) is equal to the ratio of a given dose (D) with the dose requisite for the *median* cytotoxic effect (D_M_):

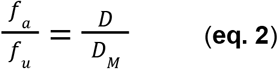

This relationship is essentially an extension of Michaelis-Menten expressions for enzyme-inhibitor equilibria ^12^. Taking the log of each side represented in **eq. 2** affords the median effect equation where log [(*f*_*a*_) / (*f*_*u*_)] is plotted as a function of log [D] to generate a linear relationship where the y-intercept is equal to -log[(D_M_)] and can be used to derive the median effect dose (D_M_). The expression in **eq. 2** is also useful for deriving additive or synergistic relationships with multiple drugs within one system. The *fraction affected* in a population of cells treated with co-administration of two inhibitors, A and B, is represented with an extension of **eq. 2** with [(*f*_*a*_)_A+B_] / [(*f*_*u*_)_A+B_] representing the affected versus unaffected fractions:

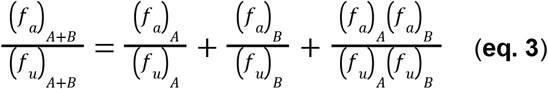

The **eq. 3** is also of high utility for representations of a **Combination Index** (**CI**) where, **CI** = [(*f*_*a*_)_A+B_] / [(*f*_*u*_)_A+B_] and the absolute value of **CI** represents the strength of non-additive behavior in the system: **CI** < 1 is synergy; **CI** = 1 is additive; **CI** > 1 is antagonistic. With **eq. 2**, it is also possible to represent the **CI** as a function of the median effect dosages:

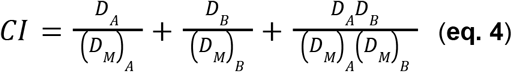

Our synergies are ranked by **CI**, and also represented with isobolograms, where the [(*f*_*a*_)_A+B_] / [(*f*_*u*_)_A+B_] is set equal to 1 (**eq. 4**). Through our analyses, we define the *fraction affected* with drug combinations for leukemia cells, across 11 distinct cell models for AML classes and subtypes including core binding factor AML, MLL-rearranged AML, and AML driven by RUNX-mutant alleles (**Table 1**).

**Table 1.**
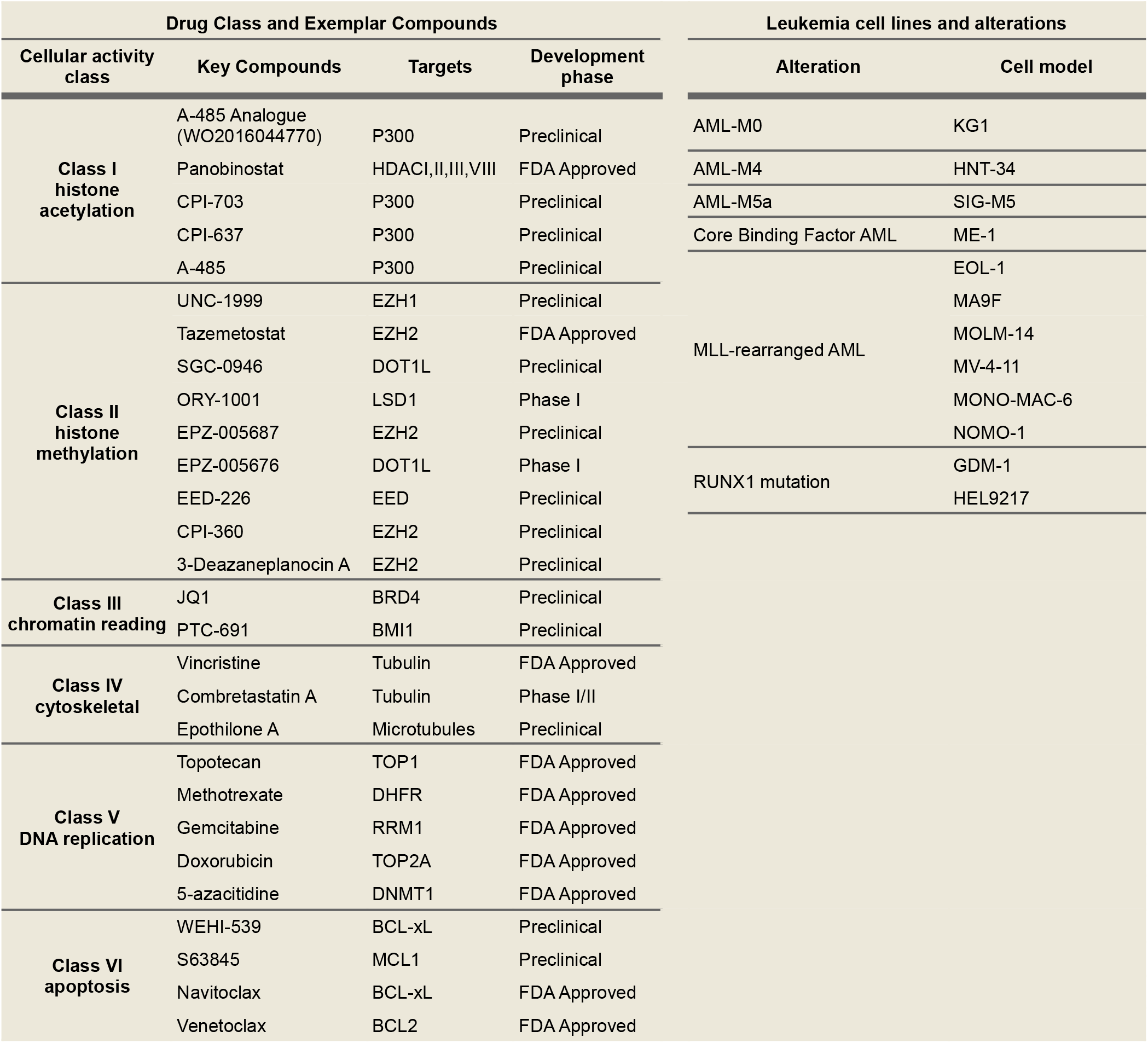
Drug classes and exemplar compounds for functional interaction studies in leukemic models.

### Synergy Analysis and Visualization

To provide our work as a public resource, we have assembled an online portal for interactive viewing of the synergy data: (https://mxw010-synergy-streamlit-example-q272y1.streamlit.app/). The code for our outward-facing resource is freely available and also open to the public (https://github.com/mxw010/synergy/blob/main/streamlit_example.py). We hypothesize that our studies will provide motivation to future pre-clinical and clinical testing of our drug combinations.

### Experimental Design

Leukemic cell lines KG1, ME-1, MOLM-14, MV-4-11, GDM1, MONO-MAC-6, HNT-34, SIGM-5, MA9F, HEL9217, and EOL-1 cells were cultured in suspension with RPMI-1640 and 20% FBS until confluency between 1-2 million cells per mL was reached. Standard conditions were used with 5% CO2 and humidity with 37 °C incubators. Upon confluency, cells were diluted to 100,000 cells per mL of culture media and distributed with multi-drop dispenser at 5 uL per well into 1536-well Corning sterile polystyrene assay plates, for a final dilution of approximately 500 cells per well. Single agent or matrix combination dosing was achieved with automated acoustic dispensing of 10 nL compounds including a proteasome inhibitor positive control and DMSO negative control within each assay plate. Combination high throughput screening was carried out with 2-fold dilutions of 9-point dose responses including controls. Edge well effects were mitigated through covering with stainless steel lids, and the plates were incubated for 48h, whereupon Cell Titer Glo (Promega) was added to each assay well in individual plates through automated liquid dispensing, and luminescence recordings were captured to quantify dose-response values ^13^.

We derive expressions for **CI** for each drug class in our synergy experiments, and represent the values in the context of isobolograms, median effects plots, and “shifts” in median effects values (IC_50_ curve shifts). Our combination studies are predicated upon rigorous single agent dose response studies, also described herein. Our data analysis pipeline is made freely available to the community and is included in the **Supplementary Methods** section.

### Open-source Automation design

In addition to the high-throughput combinatorial screening we performed, we also sought to develop an open-source implementation to enable medium-throughput screening compatible with standard laboratory liquid handling robots, which are readily available to many laboratories. Thus, the drug combination synergies we present would be amenable for replication with either system, the 1536-well automated or 384-well PyHamilton system. To establish the versatile 384-well PyHamilton system, we developed an automated combinatorial dilution robotic method, capable of performing high-n screens with a Hamilton liquid-handling robot. First, a Python library Drug Dilutions: (https://github.com/stefangolas/drug_dilutions) was generated using the PyHamilton framework (https://github.com/dgretton/pyhamilton), an open-source Python interface to Hamilton robots ^9^. Next, the main script *dispense_dilutions_2rep*.*py* parses csv files where the desired dosage pattern is represented visually as a 2-dimensional matrix corresponding to a single 384-well plate **(Figure 1A)**. The script then uses a novel algorithm to determine a time-efficient dispense pattern and executes an autonomous dispense routine, without the need to manually specify an explicit set of robot commands **(Figure 1B, Video 1)**. The ability to algorithmically determine an efficient dispense routine based on the desired dosage pattern enables flexibility across many possible dosage patterns and streamlines experimental design for combinatorial library screening. The reagents from both dose-patterned plates are then combined using the 96-channel pipetting head via quad-pinning **(Video 1)**. This method thereby enables research institutions with a variety of infrastructures to perform large numbers of combinatorial screens in-house.

**Figure 1:**
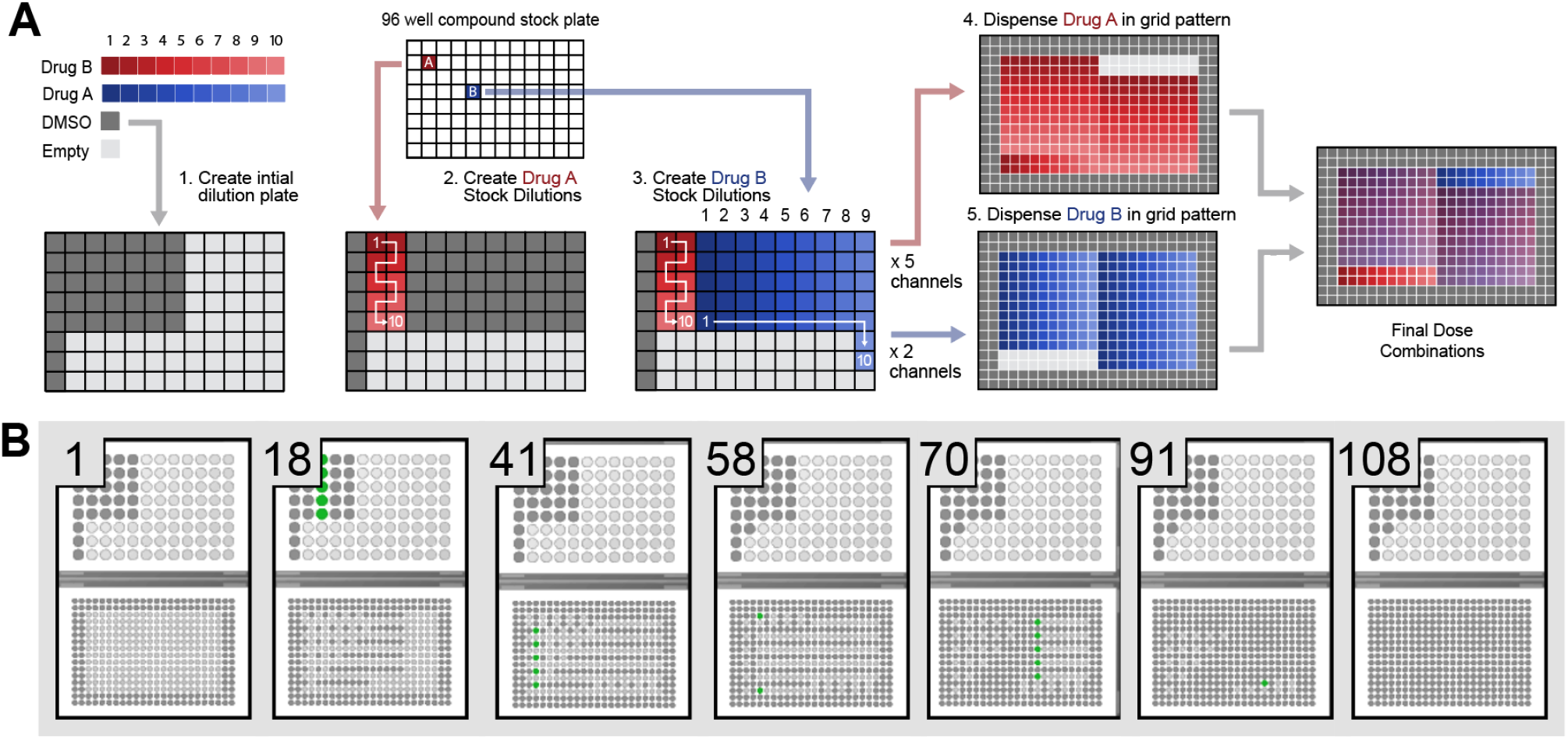
10 × 10 Dose combination performed by liquid handling robot using open-source automation package (Pyhamilton). **A)** Illustration of dispense pattern for creating a 384-well plate of 10×10 dose combinations with DMSO padding. The protocol begins with a combination of 2 drugs picked from a 96-well concentrated drug plate. Then, a plate of serial 2-fold dilutions is made with column-wise patterns arranged to match columns in the individual target plates. Finally, the two target plates are combined to create the plate of dose combinations. **B)** Selected frames output from Venus Run Control simulator (Hamilton proprietary application) in the dispensing step from serial dilutions to a target plate.

### Chromatin sequencing and analysis

Following the identification of key functional interactions, chromatin sequencing was carried out as previously reported ^14–16^. Briefly, 75-bp single-end reads from H3K27ac ChIP-seq and RAD21 ChIP-seq across treated KG1a cells were mapped to hg19 using BWA. Peaks were then called using MACS2, and peaks at ENCODE blacklisted repeat elements were excluded from downstream analysis. Within each called peak, the amount of signal was quantified using ChIP-Rx normalization strategy ^17^ with equivalent amounts of drosophila chromatin spiked in at the immunoprecipitation stage, followed by normalization to the number of reads mapping to dm3 for RRPM (Reference normalized Reads Per Million mapped reads). Heatmap plots at called peaks were generated using NGSplot, and MA plots were generated using the LSD package in R (https://rdrr.io/cran/LSD/).

## RESULTS

### Screen set-up and design

To assess the effect of clinical agents in combination, we selected eleven leukemia cell lines based on diverse driving mechanisms and genetic alterations (**Table 1**). For initial studies, we profiled a curated set of approximately 2,500 clinical, preclinical, and basic science probe compounds in high throughput miniaturized dose-response experiments, with replicates^18,19^. From the multi-well dose-response studies, we were able to select compounds with functional activity in the context of leukemia cell proliferation and to define molecular concentrations proximal to the IC_50_ values for single-agent administration. Across the leukemia cell lines in our study, we did not observe global dependencies or cell lines with acute sensitivity across compound classes (**Figure 2A**). However, MA9 and MOLM-14 cells had higher sensitivity than other cell models across the single agent 2,500 compound dose response datasets (**Figure 2B**). In the context of calculating a statistically significant synergistic combination, it is essential that each molecule sampled exhibits an independent dose response in the context of each unique cell line. As such, our method samples a wide dose range to increase versatility (spanning 10 logs), to ensure that as many dose combination synergies can be calculated as possible. Importantly, not all doses sampled can be used in the analysis, as the median effect equation only exhibits linear activity between the maximum dose that produces an *f*_*a*_ = 0, and the minimum dose that produces an *f*_*a*_ = 1 (i.e. the minimum distance between the maximum and minimum of the hill slope equation). Due to the large number of doses sampled, and the variability of each dose-response from cell line to cell line, we incorporated automated dose selection *a priori* into the analysis pipeline, to ensure that the median effect equation for each condition is only tabulated in its linear range. Still, the synergy of many drug combinations remains unquantifiable with statistical accuracy, as many independent agents have no effect on a given cell type.

**Figure 2.**
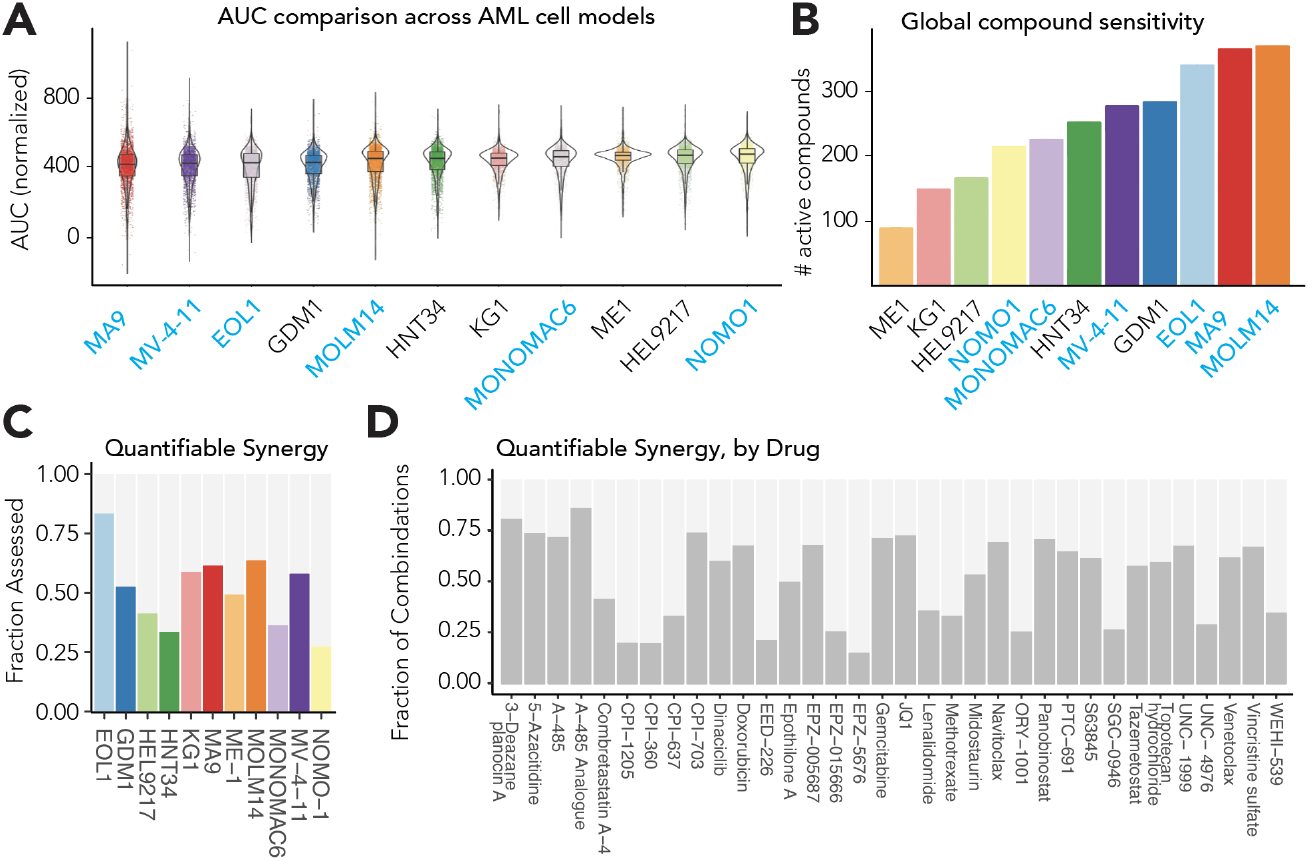
AML cell line sensitivity and synergy. **(A)** Normalized area under the dose-response curve (AUC) is plotted for MA9F, MV-4-11, EOL-1, GDM-1, MOLM-14, HNT-34, KG-1, MONOMAC-6, ME-1, HEL-2917, and NOMO-1 cells. Data are represented in violin plots to define the distribution of compound sensitivities for each AML cell line, with MLL-rearranged AML represented in blue. **(B)** Compound sensitivity is represented as the total count of active compounds in each individual AML cell line. MLL-rearranged AML is shown in blue. **(C)** The overall fraction of quantifiable synergy is represented across AML cell lines with approximately 54% synergy measurable overall. **(D)** Clinical and preclinical molecules are represented on the x-axis and the fraction of combinations with measurable synergy are represented on the y-axis.

Our initial screen was performed through the National Center for Advancing Translational Sciences (NCATS), in individual 10-point dose combinations (n=1 per dose). Collectively, of the 1300 unique drug combinations screened across 12 cell lines, 54% were able to be quantified for synergy as they contained linear median effect equations for both agents **(Figure 2C)**. However 90% of screened combinations had a reliable median effect equation for one of the two agents (with an R^2^ value > 0.7), and synergy was able to be assessed in combination with other compounds to varying degrees **(Figure 2D)**. In many cases, this is due to a lack of median-effect confidence, rather than a lack of single-agent efficacy. Thus, to further improve upon the reliability of calculating a reliable median effect equation, our open-source robotic method instead performs the 10×10 dose combination screen with an n=2 in 384 well plates. While this lowers the throughput, the increase in confidence when calculating synergy will enable researchers to readily screen hundreds of combinations of compounds without substantial data loss in future studies.

### Synergistic Drug Combinations

Interestingly, we observed high potency for Trebectidin, which is an intercalator compound that has shown strong efficacy in soft tissue sarcomas ^20^. We also observed high levels of activity also for both Daunorubicin and Doxorubicin as single-agent drugs in MV-4-11, EOL-1, GDM1, HNT34, with moderate activity in KG1, MOLM-14, HEL9217, and ME-1 (**Figure 3A**; indicated by Curve Class, **Table S1**) ^21^.

**Figure 3.**
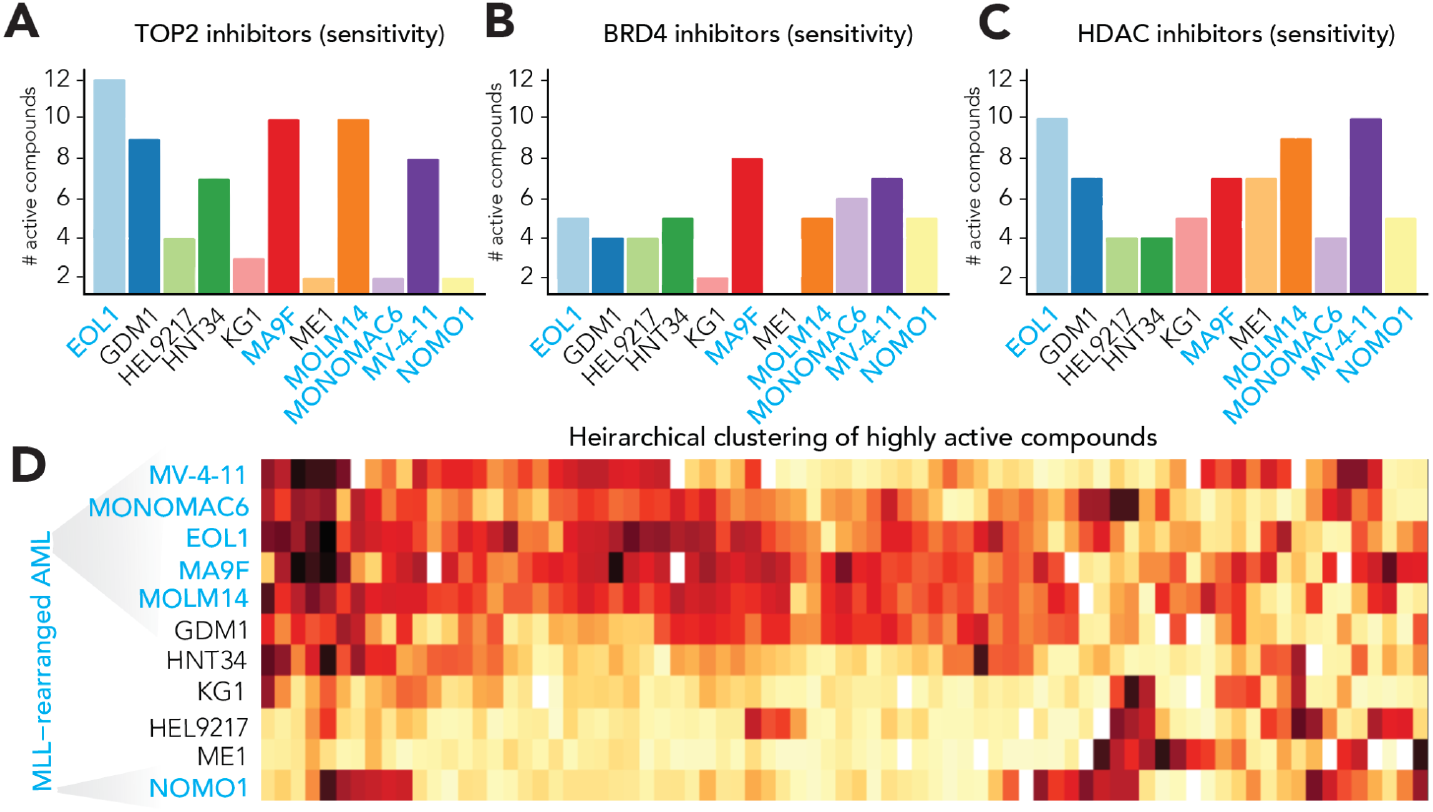
Activities in AML of molecules with single agent administration. **(A)** Selected efficacy for TOP2 inhibitors (class V), **(B)** BRD4 inhibitors (class III), and **(C)** HDAC inhibitors (class I) are represented across AML cell lines used in this study. The MLL-rearranged AML subtype is represented in blue, and the total number of active TOP2 inhibitors is on the y-axis. (**D**) Hierarchical clustering of highly active compounds with top 5% variability in the single agent screens are shown on a colorimetric scale with log_10_(AC_50_) from mM (beige) to nanomolar (red). MLL-rearranged AML is indicated in blue labeling.

These data motivated subsequent investigations of achieving increased effective potency (fraction affected) at lower D_M_ values through combination studies in non-MLL-rearranged leukemia. Across the TOPII inhibitor compound class, including anthracyclines, there was high activity in most MLL-rearranged AML cell lines and lower activity in other subtypes in our study (**Figure 3A**; **Table S1**). The trend was less clear for BRD4 inhibitors, while a similar general trend was observed for the efficacy of HDAC inhibitors in MLL-rearranged leukemia cells (**Figure 3B,C**). Further efforts will be required to understand the context-specific activities of histone deacetylase targeting in MLL-fusion driven leukemia. We observed a clustering of compound class sensitivities for MLL-rearranged AML (MONO-MAC-6, MV-4-11, EOL-1, MA9F, MOLM-14) which were highly sensitive to similar sets of compounds (**Figure 3D**; *html file 4*, heatmap 1,2). The moderate activities of drugs affecting histone H3 lysine acetylation (Class I; **Table I**), and moderate activity of anthracyclines in non-MLL rearranged AML prompted us to initiate drug synergy studies to identify compounds capable of potentiating these primary agents in these leukemia subtypes. We acknowledge the caveats that compounds from each of these six classes (**Table 1**) may have cross-talk between competing mechanisms under certain conditions, or gene expression patterns may change under the treatment of epigenetic inhibitors. We sought to understand patterns of drug-drug interactivity in AML. Thus, we operationalized high-throughput combination drug screening in 1536-well format with 5 uL volumes and acoustic dispensing of compounds ^13,19^. In our synergy studies, several key trends emerged. We noted that EZH2 inhibitors synergized strongly with anthracyclines in AML-M0, and Core binding factor AML, but did not synergize effectively in MLL-rearranged AML or AML with RUNX1-mutations (**Figure 4A-D**).

**Figure 4.**
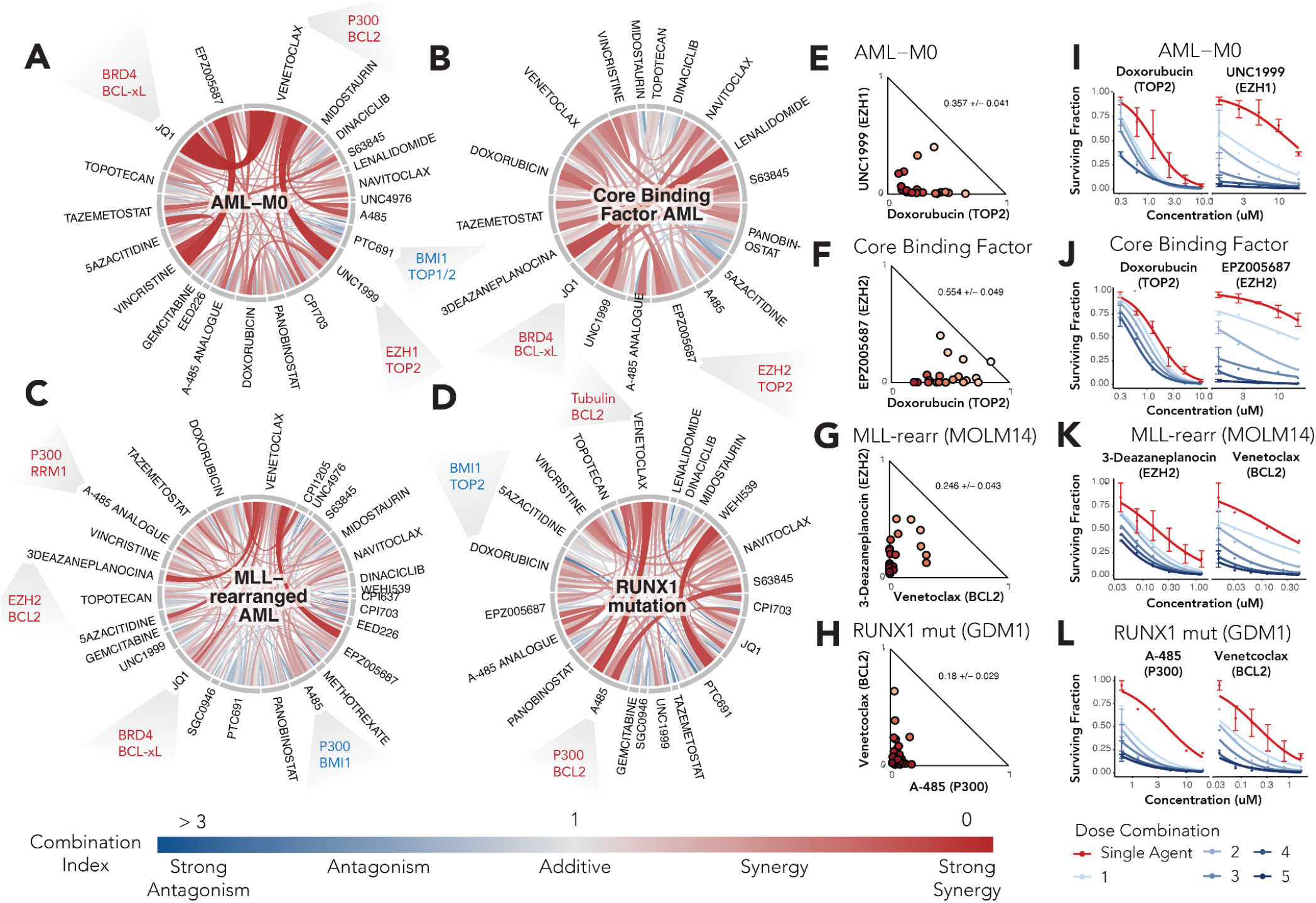
Synergy in AML subtypes with inhibitor combinations. Circos plots represent drug synergies in **(A)** AML-M0, **(B)** Core Binding Factor AML, **(C)** MLL-rearranged AML, and **(D)** RUNX1-mutant AML. In each case, the intensity (red) of the arc denoting drug-drug interactions is a surrogate for the strength of the synergy, while blue denotes functional antagonism with drug-drug interaction pairs. Isobolograms are shown for **(E)** AML-M0, **(F)** Core Binding Factor AML, **(G)** MLL-rearranged AML, and **(H)** RUNX1-mutant AML, with highly representative drug combinations shown for each subtype. Combinations are shown for inhibitors that were tested with at least 3 other molecules. Synergies affecting the placement of modifications on histone H3 lysine-27 are represented in dose-response curve shift plots for **(I)** AML-M0, TOP2-EZH1 **(J)** Core Binding Factor AML, TOP2-EZH2 **(K)** MLL-rearranged AML, EZH2-BCL2 and **(L)** RUNX1-mutant AML, A485-BCL2. In each case, one drug shows minimal single-agent activity in the AML cells, but is strongly potentiated by the second agent.

It was particularly notable that chromatin depressing drugs (e.g., Tazemetostat, EPZ005687, UNC1999) had strong synergistic relationships with doxorubicin, which may require accessible DNA to function ^22,23^. We emphasize the importance of EZH2 synergizers, given the relatively non-toxic effects of EZH2-targeting compounds within 48h, while sub-IC50 level co-administration with anthracyclines elicited strongly toxic effects with clinical EZH2 drugs like tazemetostat. Further work will be required to understand the precise mechanisms for how chromatin derepression potentiates anthracyclines in AML. We generated isobolograms for UNC1999-Doxorubicin (**Figure 4E**) and EPZ005687-Doxorubicin (**Figure 4F**) and noted the strength and breadth of the EZH2-anthracycline synergies, where many doses exhibited strong synergism.

In MLL-rearranged AML particularly strong synergy was observed with Venetoclax and 3-Deazaneplanocin (DZNep), which represented drug class synergy between EZH2 and BCL2 (Class II, Class VI, **Table I**; **Figure 4G,K**). This may present new opportunities for developing clinical drug combinations with FDA-approved BCL2 inhibitors in future studies. Other examples of Class II drug synergies with Class VI were the A484-Venetoclax drug interactions we observed in RUNX1-mutant AML (**Figure 4H,L**). We also observed strong antagonisms in our datasets. The BMI1 inhibitor PTC-691 was strongly antagonistic with anthracyclines across AML subtypes (**Figure 4A,C,D**). This was intriguing mechanistically, because of the expectation that EZH2-anthracycline synergy should predict BMI1 functional synergy as well. However, we noted that PTC-691 is known to inhibit the transcription of BMI1, or stability of its post-translational modifications, in specific cellular contexts, which may introduce alternative mechanistic explanations for potential restriction of chromatin sites for Doxorubicin in the presence of PTC-691 (**Table 1**). Taken together, we observed strong shifts in IC50 values for EZH1/2-anthracycline synergies (**Figure 4E,G**), and BCL2 synergies with drugs affecting the modification state on histone H3 Lys-27 (**Figure 4I,K**).

### Genomics of Drug Synergy

In addition to the synergies between Class II and Class V (EZH2-Anthracycline), or Class II and Class VI (EZH2-BCL2), we also observed unexpected intra-Class II synergy. In replicate experiments, we observed EZH2 inhibitor EPZ005687 synergy with the P300 inhibitor A-485 across a range of concentrations (**Figure 5A**). The novelty of molecular synergy affecting post-translational modifications on the same histone tail, H3 Lys-27 motivated us to investigate further to examine the functional consequences of the drug combination on chromatin. We examined chromatin immunoprecipitation coupled with next-generation sequencing ^14,24–26^. In our studies, we first sought to “digitize” the readout for H3K27ac losses with A484 in AML. Thus, we performed H3K27ac ChIP-seq within hours of A485 treatment and uncovered thousands of lost H3K27ac signal “peaks” in the data (**Figure 5B,C,D**). However, when examining synergistic losses of H3K27ac with combined treatment with EZH2 inhibition and P300 inhibition, we observed a surprising enhancement of H3K27ac losses across the genome, with newly formed H3K27ac peaks only representing a small subset of the data (**Figure 5B,C,D**). For these studies, we selected the 4h timepoint to avoid global non-specific effects of cell cycle exit concomitant with toxic drug treatments over longer periods.

**Figure 5.**
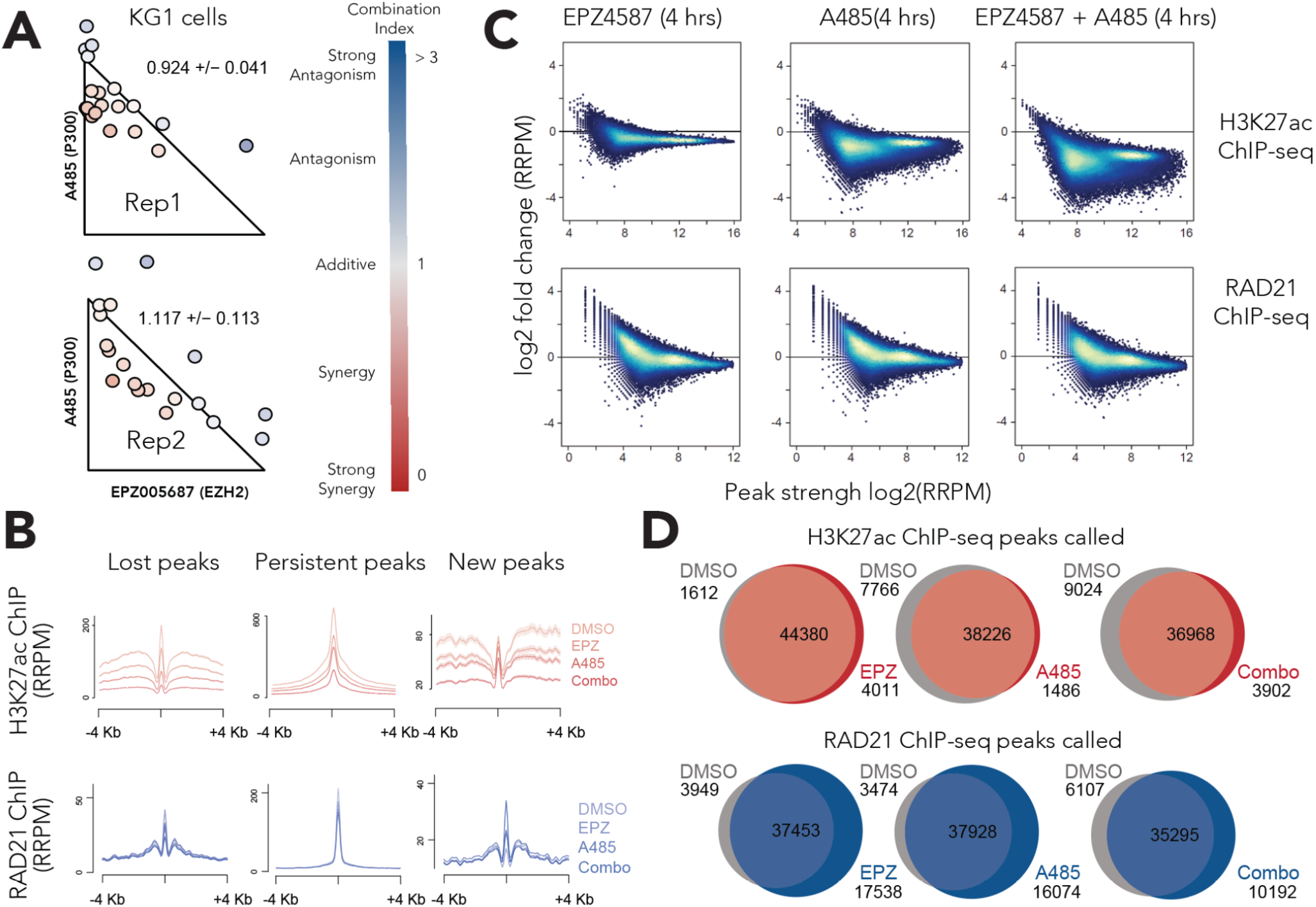
Mechanistic epigenetics from functional synergy in AML. **(A)** Isobolograms for A485-EPZ005687 (P300-EZH2 synergy) are represented for KG1 cells, with two replicates. The colorimetric index illustrates the strength of synergy or antagonism in red or blue, respectively. **(B)** Mechanistic evaluation of the EZH2-P300 synergy is shown with ChIP-seq for H3K27ac (top) or Cohesin complexes (RAD21, bottom) with DMSO, EPZ005687, A485, or EPZ005687-A485 (combo) from lightest to darkest intensity scale in red (H3K27ac) or blue (RAD21). The drug combinations produce lost binding sites for Cohesin (bottom left) and lost deposition of H3K27ac (top left) suggesting that drug synergy also deregulates chromatin structure synergistically. **(C)** MAPlots of rapid (4h) treatments of EZH2 drug (EPZ005687), P300 drug (A485), or EZH2-P300 combination (EPZ5687 + A485) are shown in the context of genome-scale changes in placement of H3K27ac (top) or bottom (RAD21). Each datum point is represented in the MAPlots by blue circles, with ChIP-seq peak intensity on the x-axis and log2 fold change RPM values on the y-axis. (**D**) Total ChIP-seq peak counts are listed with venn diagrams showing overlaps between control and treatment conditions for DMSO-vs-EPZ005697 (EPZ), DMSO-vs-A485, or DMSO-vs-EPZ005687 + A485 (combo). H3K27ac peaks are shown (top) and RAD21 peaks are shown (bottom) for each treatment set.

We next examined the effects of drug synergy on the placement of Cohesin complexes, which represent an essential layer to folding the 3D genome ^27,28^. We observed a unique effect in the context of RAD21 ChIPseq (RAD21 is a key subunit of mammalian Cohesin complexes), where Cohesin actually gained thousands of peaks across the genome with EZH2 inhibition (**Figure 5D**). Whether these data represent an additional regulatory relationship between PcG complexes and Cohesin will require further investigation. We next sought to determine whether combination P300/EZH2 drug treatments elicited similar effects on Cohesin localization as on deposition of the H3K27ac mark. We observed that global loss of Cohesin binding with combined drug treatments was not nearly as profound as for placement of the H3K27ac mark (**Figure 5C,D**), while we did note that under combined drug treatment conditions Cohesin migrated to many new or unique genomic binding sites (**Figure 5B)**. Further work will reveal whether the deregulation of Cohesin function represents a vulnerability in leukemia, while genetic evidence points towards RAD21 and the SMC1/3 proteins as being essential in this tumor ^28^.

## Discussion

While tumor heterogeneity and drug resistance have each become major themes in cancer research, we are only beginning to understand the significance of combinatorial drug therapy in this context. There is increasing evidence that lethal tumors can evolve rapidly to compensate for single-agent drug treatment ^29–32^. Understanding the mechanisms of efficacious drug combinations, and the generation of sufficiently large and statistically reliable datasets will advance the larger goal of rational prediction of which classes of drugs may work in a particular genetic or disease context. In our studies, we have examined thousands of single-agent drugs for efficacy in AML subtypes, and hundreds of combinatorial drug combinations in high throughput, and in functional chromatin sequencing assays.

Our studies have revealed that drugs affecting the modification states for lys-27 on histone H3 can synergize effectively with drugs affecting cell cycle (Class V), regulation of apoptosis (Class VI), in particular. Of note, our unbiased high-throughput approaches have revealed that unexpected combinations of inhibitors each targeting opposing chromatin modification states on histone H3 Lys-27 can synergize and alter the deposition of H3K27ac across the genome in AML. Our work also generalizes opportunities for high throughput combination drug discovery with open-sourced automation and computational pipelines and provides a large new reliable dataset for current or future machine-guided learning efforts. Remaining questions from these efforts include understanding how anthracyclines potentiate EZH2 inhibition at the chromatin level, which will be of high interest ^33^. Together, this resource will allow labs to generate combinatorial synergies between drugs, to catalyze our efforts as a community to address resistance and tumor evolution.

Given the high degree of efficacy for the “CHOP” drug combination regimen for lymphomas (including cyclophosphamide, doxorubicin hydrochloride (hydroxydaunorubicin), vincristine sulfate (Oncovin), and prednisone), we hypothesize that discovery of new combination therapies for leukemias will be catalyzed by rigorous preclinical drug combination studies, and efficacy rankings (see **Table S1**). It is of note that certain combination synergies herein we could not have predicted *a priori*, including those where epigenetic drugs target opposing modifications on the same histone tail (e.g., Panobinostat, A485). Thus, we believe that empirically driven synergy discovery will be impactful for drug repurposing, and for identifying unexpected pathways through which drugs may act in combination, and predicting new and uncharted drug combinations that may ultimately improve patient outcomes.

## Acknowledgments

We are grateful to Drs. B. Mizukawa, M. Wunderlich E. O’Brien, T. Guinipero, S. Michael, K. Wilson, P. Shin, M. Hall, A. Simeonov, M. Ferrer, D.Y. Duveau, C.E. Jones, R.D. Roberts, and H. Liu for helpful comments and assistance. Special thanks to the Andrew McDonough B+ Foundation for supporting this work, and other projects to define new therapies and basic advances for leukemia.

